# Recombinant Rotavirus Expressing the Glycosylated S1 Protein of SARS-CoV-2

**DOI:** 10.1101/2023.08.01.551500

**Authors:** Asha A. Philip, Sannoong Hu, John T. Patton

**Affiliations:** Department of Biology, Indiana University, Bloomington, IN 47405, USA

**Keywords:** rotavirus, expression vector, SARS-CoV-2, glycosylation, spike protein

## Abstract

Reverse genetic systems have been used to introduce heterologous sequences into the rotavirus segmented double-stranded (ds)RNA genome, enabling the generation of recombinant viruses that express foreign proteins and possibly serve as vaccine vectors. Notably, insertion of SARS-CoV-2 sequences into the segment 7 (NSP3) RNA of simian SA11 rotavirus was previously shown to result in the production of recombinant viruses that efficiently expressed the N-terminal domain (NTD) and the receptor-binding domain (RBD) of the S1 region of the SARS-CoV-2 spike protein. However, efforts to generate a similar recombinant (r) SA11 virus that efficiently expressed full-length S1 were less successful. In this study, we describe modifications to the S1-coding cassette inserted in the segment 7 RNA that allowed recovery of second-generation rSA11 viruses that efficiently expressed the ∼120-kDa S1 protein. The ∼120-kDa S1 products were shown to be glycosylated, based on treatment with endoglycosidase H, which reduced the protein to a size of ∼80 kDa. Co-pulldown assays demonstrated that the ∼120-kDa S1 proteins had affinity for the human ACE2 receptor. Although all the second-generation rSA11 viruses expressed glycosylated S1 with affinity for the ACE receptor, only the S1 product of one virus (rSA11/S1f) was appropriately recognized by anti-S1 antibody, suggesting the rSA11/S1f virus expressed an authentic form of S1. Probably due to the presence of FLAG tags on their S1 signal peptides, the S1 products of the other viruses (rSA11/3fS1 and rSA11/3fS1-His) may have undergone defective glycosylation, impeding antibody binding. In summary, these results indicate that recombinant rotaviruses can serve as expression vectors of foreign glycosylated proteins, raising the possibility of generating rotavirus-based vaccines that can induce protective immune responses against enteric and mucosal viruses with glycosylated capsid components, including SARS-CoV-2.

## Introduction

Rotaviruses are a large genetically diverse group of segmented double-stranded (ds) RNA viruses that belong to the family *Sedoreoviridae* (1, 2). The viruses are important human pathogens, representing the primary cause of acute potentially life-threatening gastroenteritis in children under 5 years of age (3, 4). The extensive morbidity and mortality associated with rotavirus infections have stimulated the development of rotavirus vaccines and their introduction into the childhood immunization programs of many countries, including the US (5, 6). The most widely used rotavirus vaccines are formulated from live attenuated strains of the virus and are given orally to infants during the first 6 months of life (7).

The rotavirus virion is a non-enveloped triple-layered icosahedron particle that encapsidates 11 segments of dsRNA (8). The genome segments of prototypic strains of group A rotaviruses (Rotavirus A species (RVA)) have a total size of ∼18.5 K base pairs (Kbp) and encode 6 structural (VP1-VP4, VP6-VP7) and 6 nonstructural proteins (NSP1-NSP6) (2, 9). Development and refinement of plasmid-based rotavirus reverse genetics systems have allowed the generation of recombinant RVA viruses of diverse origin, including strains isolated from humans, non-human primates, and rodents (10–13). Rotavirus reverse genetics systems have been used to manipulate the protein products encoded by most of the genome segments of the virus (14–21).

Several studies have described the isolation and characterization of rotavirus variants that, due to sequence duplications, contain one or more unusually large genome segments (22–26). The most commonly described variants are those with sequence duplications present within segment 5 (27), which encodes the viral interferon antagonist NSP1 (28, 29), or segment 7, which encodes the viral translation enhancer NSP3 (30–32). Variants have been identified with sequence duplications in segment 5 of almost 1.5 Kbp, attesting to the capacity of the rotavirus genome and virus particle to accommodate significant amounts of extra sequence (27).

Recombinant rotaviruses have been made by reverse genetics that, through the introduction of heterologous sequences into segment 5 and 7 RNAs, are able to express foreign proteins (33). Because NSP1 is not essential for rotavirus replication, some recombinant rotaviruses expressing foreign proteins have been made by replacing portions of the NSP1 ORF with heterologous coding sequences (34–36). Other recombinant viruses have been made by placing the heterologous coding sequence at the end of the full-length NSP1 ORF (37).

Recombinant rotaviruses that express foreign proteins from segment 7 have been generated though placement of heterologous coding sequences at the end of the NSP3 ORF (38–40). Such segment 7 modifications have been used to prepare recombinant viruses that express NSP3 fusion proteins, for example, NSP3 fused to the green epifluorescence protein UnaG (NSP3-UnaG) (39). By insertion of a 2A stop-restart translation element in segment 7, between the coding sequence for NSP3 and a foreign protein, recombinant rotaviruses have been generated that express two separate protein products from segment 7 (38, 40, 41). This approach has allowed the development of rotaviruses that express the complete complement of rotavirus proteins plus an additional non-rotaviral protein (40). Recombinant rotaviruses that express immunogenic regions of the capsid proteins other viruses may allow for the generation of combination vaccines that can induce protective immune responses to not only rotavirus, but also a second pathogenic virus, such as norovirus (42).

In earlier studies, we investigated the possibility of making recombinant rotaviruses that expressed portions of the spike (S) protein of SARS-CoV-2 (43). In this work, recombinant SA11 (rSA11) rotaviruses with segment 7 modifications were recovered that expressed the N-terminal domain (NTD), the receptor binding domain (RBD), and the core domain (CR) of the SARS-CoV-2 S protein (44, 45). A similar segment 7 modification was used to make a recombinant virus (rSA11/fS1) containing the complete coding sequence of the SARS-CoV-2 S1 protein, a cleavage fragment of the S protein that includes both the NTD and RBD and is a primary target of neutralizing antibodies produced during SARS-CoV-2 infection (46–50). The open reading frame (ORF) in the modified segment 7 RNA of the rSA11/fS1 virus included the coding cassette NSP3-2A-3xFLAG-S1. Through the action of the 2A translation element, the segment 7 RNA of the virus was expected to generate two products: NSP3 fused to a 2A peptide (NSP3-2A) and 3xFLAG-tagged S1 (fS1). In the NSP3-2A-3xFLAG-S1 cassette, a 3xFLAG tag was positioned immediately upstream of the S1 signal peptide, an element critical for synthesis of glycosylated S products (51). Immunoblot analysis of the products made by rSA11/fS1 indicated that although the virus efficiently made NSP3-2A, it was not efficient in generating the expected fS1 product, possibly due to instability or degradation of the S1 product, or impact of the FLAG tag on the function of the signal peptide (43). In the work described here, we have compared the S1 products made by the rSA11/fS1 virus to the products made by newly designed rSA11 viruses encoding S1 proteins differing in the nature of their terminal peptide tags. The results showed that one of newly designed rSA11 viruses (rSA11/S1f) efficiently expressed S1 protein and that the S1 protein were glycosylated, localized to ER/Golgi vesicles, had affinity for the extracellular domain of the ACE2 receptor (52). This is the first study demonstrating that recombinant rotaviruses can be used as expression vectors of glycosylated foreign proteins.

## Materials and methods

### Cell culture

Embryonic monkey kidney cells (MA104) were grown in Dulbecco’s modified Eagle’s medium (DMEM) containing 4.5 g/L glucose (Lonza 12-640F or Corning 15-107-CV), 1% penicillin-streptomycin [Corning]), and 5% fetal bovine serum (FBS, Gibco) (Arnold et al, 2009). Baby hamster kidney cells constitutively expressing T7 RNA polymerase (BHK-T7 cells) were kindly provided by Drs. Ulla Buchholz, Laboratory of Infectious Diseases, NIAID, NIH. BHK-T7 cells were grown in Glasgow complete medium (GMEM, Lonza) supplemented with 10% tryptone-peptide broth (Gibco), 1% penicillin-streptomycin Gibco), 2% non-essential amino acids (Gibco), 1% glutamine (Gibco), and 5% heat-inactivated FBS (53). Medium used to cultivate BHK-T7 cells was supplemented with 2% G418 (Geneticin, ThermoFisher) every other passage.

### Plasmids

Plasmids used in generating recombinant SA11 rotaviruses were obtained from Addgene [https://www.addgene.org/Takeshi_Kobayashi/] and included pT7/VP1SA11, pT7/VP2SA11, pT7/VP3SA11, pT7/VP4SA11, pT7/VP6SA11, pT7/VP7SA11, pT7/NSP1SA11, pT7/NSP2SA11, pT7/NSP3SA11, pT7/NSP4SA11, and pT7/NSP5SA11. The plasmids pCMV-NP868R, pT7/NSP3-P2A-fUnaG, and pTWIST/COVID19spike were derived as described earlier (39, 40, 43). The plasmid pT7/NSP3-2A-3fS1 was generated as described by Philip and Patton (43) and contains a full-length cDNA of the SARS-CoV-2 spike S1 open reading frame (ORF) (GenBank MN908947.3). The plasmids pT7/NSP3-2A-S1f and pT7/NSP3-2A-3fS1-His contain the same S1 ORF as pT7/NSP3-2A-3fS1 but differ in the nature of sequences for peptide tags surrounding the S1 ORF.

The pT7/NSP3-2A-S1f plasmid was constructed using a Takara In-Fusion cloning kit, which combined the vector backbone (pT7/NSP3-P2A region) of pT7/NSP3-P2A-fUnaG (primer pair for amplification: TGACCATTTTGATACATGTTGAACAATCAAATACAG and AGGACCGGGGTTTTCTTCCAC) with the S1 ORF insert of pTWIST/COVID19spike (primer pair: GAAAACCCCGGTCCTGTGTTTGTTTTTCTTGTTTTATTGCCACTAGTCT and GTATCAAAATGGT<u>CACTTGTCATCGTCATCCTTGTAATC</u>ACGTGCCCGCCG). Primers were designed to include a sequence for a 1X FLAG tag at the C-terminus of the encoded S1 protein (underlined). The pT7/NSP3-2A-3fS1-His plasmid was produced by inserting a sequence encoding a 6X His tag at the 3’-end of the S1 ORF in pT7/NSP3-2A-3fS using an In-Fusion cloning kit. This was accomplished by amplifying pT7/NSP3-2A-3fS with the primer pair: ACCACCACCACCACCACTGACCATTTTGATACATGTTGAACA and GGTGGTGGTGGTGGTGACGTGCCCGCCGAGGAGA. Transfection quality plasmids were prepared using Qiagen plasmid purification kits. Primers were obtained from Eurofins Scientific and plasmid sequences were verified by Eurofins Genomics.

### Recombinant viruses

Detailed procedures for generating and recovering recombinant SA11 rotaviruses have been published before (39, 53). Briefly, BHK-T7 cells were transfected with SA11 pT7 plasmids and pCMV-NP868R using Mirus TransIT-LT1 transfection reagent. pT7/NSP2SA11 and pT7/NSP5SA11 were included in transfection mixtures at levels 3-fold higher than the other plasmids. As necessary, the pT7/NSP3SA11 plasmid was replaced with pT7/NSP3-2A-3fS, pT7/NSP3-2A-S1f or pT7/NSP3-2A-3fS1-His. The transfected cells BHK-T7 cells were overseeded with MA104 cells at 2 days post infection, and the growth medium was adjusted to a final concentration of 0.5 μg/ml trypsin (porcine Type IX pancreatic trypsin, Sigma Aldrich). Once complete cytopathic effects (CPE) were observed, cells in the media overlay were subject to three rounds of free-thaw and the lysate clarified by low speed centrifugation. Virus in lysates were recovered by plaque isolation and amplified by one round of growth on MA104 cells (53, 54). Viral dsRNAs were recovered by TRIzol (Thermo Fisher) extraction, resolved by polyacrylamide gel electrophoresis, and detected by staining with ethidium bromide (53, 54). cDNAs were generated from dsRNAs using a Superscript III One-Step RT-PCR Platinum Taq kit (Thermo Fisher) and appropriate segment 7 (NSP3) primers and sequenced by Eurofins Genomics.

### Immunoblot analysis

Proteins present in MA104 cell lysates were detected by immunoblot assay following previously described procedures (42, 43). Cells were mock infected or infected with 5 plaque forming units (PFU) of recombinant virus, collected at 9 h p.i., and lysed by resuspending in immunoprecipitation (IP) lysis buffer (300 mM NaCl, 100 mM Tris-HCl, pH 7.4, 2% Triton X-100) containing ethylenediaminetetraacetic acid (EDTA)-free protease inhibitor cocktail (Roche cOmplete, Sigma Aldrich)]. Proteins were resolved by electrophoresis on 10% polyacrylamide (SDS) gels and transferred to nitrocellulose membranes using a Bio-Rad Trans-Blot Turbo Transfer System. Membranes were blocked with phosphate-buffered saline containing 5% non-fat dry milk and probed with rabbit polyclonal SARS-CoV-2 S1 antibody (A20136, ABclonal, 1:1000 dilution), guinea pig polyclonal NSP3 (NIH Lot 55068, 1:2000 dilution) or VP6 (NIH Lot 53963, 1:2000) antisera, mouse monoclonal FLAG M2 (F1804, Sigma-Aldrich, 1:2000) or His-tag antibody (MCA1396, Bio-Rad, 1:1000), or rabbit monoclonal β-actin antibody (D6A8, Cell Signaling Technology, 1:1000). In some cases, blots were reprobed with a different antibody following treatment with WesternSure ECL stripping buffer (LI-COR Biosciences). Primary antibodies were detected using 1:10,000 dilutions of horseradish peroxidase (HRP)-conjugated secondary antibodies: horse anti-mouse IgG (Cell Signal Technology), goat anti-guinea pig IgG [Kirkegaard & Perry Laboratories (KPL), or goat anti-rabbit IgG (Cell Signaling Technology). Signals were developed using Clarity Western ECL Substrate (Bio-Rad) and detected using a Bio-Rad ChemiDoc imaging system.

### Endoglycosidase H (Endo H) assay

MA104 cell monolayers in 6-well plates were mock-infected or infected with rSA11 viruses (5 PFU per cell). At 9 h p.i., cell monolayers were washed and scraped into phosphate-buffered saline (PBS), pelleted by low-speed centrifugation, and resuspended in 250 μl per well of IP lysis buffer. The presence of glycosylated proteins in the cell lysates was assessed using Promega Endoglycosidase H (Endo H) assay reagents (Promega V4871). Briefly, 27 μl samples of cell lysates were combined with 3 μl of 10X Denaturing Solution, heated to 95°C for 5 min, and cooled to room temperature. The heat-treated lysates were mixed with 3 μl of nuclease-free water, 4 μl of 10X Endo H Reaction Buffer and 3 μl of Endo H enzyme, then incubated at 37°C for 16 h. Proteins in the processed samples were detected by immunoblot assay, as described above.

### S1-ACE2 interaction assay

A Takara Capturem IP and Co-IP kit (Cat No: 635721) was used to assess the affinity of SARS-CoV-2 S1 expressed by rSA11 viruses for ACE2. Protein A spin columns and all necessary buffers were included in the Capturem IP kit. MA104 cell monolayers were mock-infected or infected with rSA11 viruses (5 PFU/cell). At 9 h p.i., the cells were washed and scraped into PBS, pelleted by low-speed centrifugation, and resuspended in Lysis/Equilibration Buffer containing protease inhibitor cocktail. After a 15 min incubation on ice, the lysate was clarified by centrifugation at 17,000 *g* for 10 min. Soluble hACE2-Fc (fchace2, InvivoGen), a recombinant protein consisting of the extracellular domain of human ACE2 fused to a human IgG1 Fc region, was added to the clarified lysates, to a final concentration of 20 μg per ml, and the mixture incubated overnight at 4°C. To recover complexes formed between the hACE2-Fc and S1 proteins, lysate samples were loaded onto pre-equilibrated protein A spin columns, which were then centrifuged at 1000 *g* for 1 min at room temperature. After rinsing columns with Wash Buffer, proteins were eluted from columns by adding Elution Buffer and centrifugation at 1000 *g* for 1 min at room temperature. The eluted samples were immediately neutralized by adding Neutralization Buffer. Proteins in eluted samples were detected by immunoblot assay, as described above.

### Immunofluorescence analysis

MA104 cells were grown on poly-L-lysine-coated coverslips in 12-well culture dishes and mock infected or infected with 5 PFU of trypsin-activated rSA11 virus per cell. Prior to infection, virus inoculum was trypsin-activated by incubating with 10 μg porcine trypsin, type IX-S (Sigma-Aldrich), per ml for 30 min at 37°C. At 9 h p.i., the cells were fixed by incubation with ice cold methanol for 3 min, then washed twice with PBS containing 1% Triton X-100 (PBS-TX). The cells were incubated in PBS containing 5% bovine serum albumin (BSA) for 30 min at room temperature and then incubated with rabbit polyclonal S1 antibody (ABclonal A20136), and mouse monoclonal NSP2 (lot #171, 1:1500) or NSP4 antibody (lot #55/4, 1:800) in PBS containing 3% BSA for 1 h at room temperature. Afterwards, the cells were washed thrice with PBS-TX, followed by incubation with Alexa 488 anti-rabbit IgG (green) and Alexa 594 anti-mouse IgG (red) (Molecular Probes) in PBS containing 3% BSA for 30 min at room temperature. The cells were washed 3-times with PBS-TX, and the coverslips were mounted with ProLong antifade reagent containing 4,6-diamino-2-phenylindole (DAPI) (Invitrogen). Cells were analyzed with a Nikon Eclipse NiE microscope (100X oil immersion objective) and images were captured with a Hamamatsu Orca-Flash 2.8 sCMOS high resolution camera using GFP and Alexa 594 windows.

### Genetic stability

The genetic stability of recombinant rotaviruses was assessed by serial passage on MA104-cell monolayers using 1:10 dilutions of infected cell lysates prepared in serum-free DMEM and 0.5 µg/ml trypsin (42, 54). Viral dsRNA was recovered by Trizol extraction from clarified cell lysates treated with RNase T1 to remove single-stranded RNA (39, 53). Viral dsRNA was analyzed by electrophoresis on 8% polyacrylamide gels and detected by straining with ethidium bromide.

### GenBank accession numbers

Modified segment 7 sequences of rSA11 viruses that have been deposited in GenBank: rSA11/wt (LC178572), rSA11/3fS1 (MW059026), rSA11/S1f (MZ511690), and rSA11/3fS1-His (MZ511689). Other accession numbers include the SARS-CoV-2 S sequence in pTWIST/COVID19spike (GenBank MN908947), sequence for the African swine fever virus capping enzyme in pCMV-NP868R, and modified segment 7 RNA of rSA11-NSP3-P2A-3fUnaG (MK851042). This study was approved by Indiana University under IBC Protocol BL-879-13.

## Results

### Generation of recombinant viruses encoding S1 protein

In a previous study, a rSA11 virus (rSA11/3fS1) was generated with a modified segment 7 (NSP3) RNA that encoded the SARS-CoV-2 S1 protein with a fused N-terminal 3xFLAG tag (NSP3-2A-3fS1) (Fig. 1). Notably, the 3xFLAG tag was positioned immediately upstream of the N-terminal signal peptide of the S1 protein. To gain a better understanding of factors affecting the poor expression of S1 by rSA11/3fS1, we generated two similar rSA11 viruses differing only in the nature of the peptide tags encoded upstream and downstream of the S1 ORF in the segment 7 RNA. One of the viruses, rSA11/3fS1-His (Fig. 1), was identical to rSA11/NSP3-2A-3fS1, with the exception that the ORF in its segment 7 RNA was engineered to introduce a 6xHis tag at the end of the S1 product. The rSA11/3fS1-His virus was generated to address the possibility that, due to cleavage of the signal peptide from the S1 product, the N-terminal 3xFLAG tag was lost, preventing accurate assessment of fS1 synthesis by the rSA11/NSP3-2A-3fS1 virus via immunoblot assay with anti-FLAG antibody (43). Instead, the production of S1 products could be assessed with anti-His antibody. The second recombinant virus that was made, rSA11/S1f (Fig. 1), contained a segment 7 RNA designed to express S1 with a C-terminal 1xFLAG tag, but without any N-terminal tag (NSP3-2A-S1f). The usefulness of this virus was in examining the possibility that a tag positioned upstream of the S1 signal peptide might impede synthesis and glycosylation of the S1 protein on the endoplasmic reticulum.

**Figure 1.**
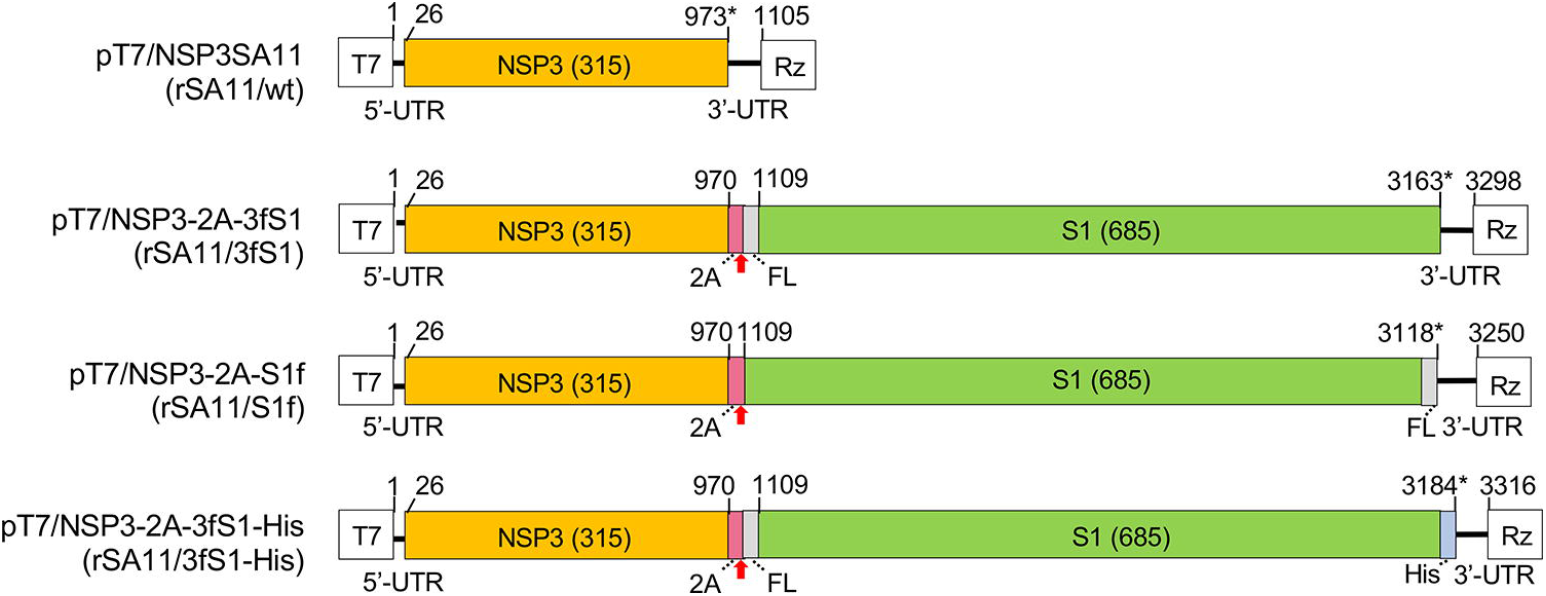
Modified segment 7 (NSP3) plasmids used to generate rSA11s encoding the SARS-CoV-2 S1 protein. Schematic indicates the nucleotide positions of the coding sequences for NSP3, porcine teschovirus 2A element, 3X or 1X FLAG (FL), 6X His (His) and the complete S1 in pT7 plasmids. The red arrow notes the position of the 2A translational stop-restart site, and the asterisk notes the end of the ORF. Sizes (aa) of encoded NSP3 and S1 proteins are given Virus (rSA11) recovered using the pT7 plasmid is indicated in parenthesis. T7 (T7 RNA polymerase promoter sequence), Rz (Hepatitis D virus ribozyme), UTR (untranslated region).

The recombinant viruses, rSA11/S1f and rSA11/3fS1-His, were produced following the same reverse genetics procedure used previously to generate rSA11/3fS1 and the wildtype virus, rSA11/wt (39, 43). The procedure included transfection of BHK-T7 cells with a set of T7 transcription vectors (pT7) expressing SA11 plus-sense (+)RNAs and a CMV expression plasmid (pCMV-NP868R) encoding the capping enzyme of African swine fever virus. In the transfection mixtures, T7 transcription vectors for NSP2 and NSP5 +RNAs (pT7/SA11NSP2 and pT7/SA11NSP5, respectively) were used as levels 3-fold greater that the other pT7 vectors. The modified segment 7 transcription vectors (pT7/3fS1, pT73fS1-His, and pT7/S1f, Fig. 1) used in generating rSA11 encoding S1 products was added to transfection mixtures in place of pT7/NSP3SA11. Recombinant viruses formed in transfected BHK-T7 cells were amplified by overseeding with MA104 cells and then isolated by plaque purification. Recombinant viruses were further amplified in MA104 cells.

### Genomes and growth characteristics of rSA11s

The dsRNA genome segments of recombinant rotaviruses were resolved by gel electrophoresis to verify the presence of modified segment 7 RNAs (Fig. 2A). The analysis showed that rSA11/3fS1, rSA11/S1f and rSA11/3fS1-His all lacked the 1.1-Kbp segment 7 dsRNA typical of rSA11/wt. Instead, the S1-encoding rSA11 viruses contained segment 7 dsRNAs that migrated on polyacrylamide gels near the position of the segment 1 dsRNA and had a size close to 3.3 Kbp. Thus, the segment 7 RNAs of the mutant viruses were approximately 3-times the size of SA11/wt segment 7 RNA. Sequencing verified that the segment 7 sequence of the recombinant viruses matched those present in the pT7/NSP3-2A-3fS1, pT7/NSP3-2A-3fS1-His, and pT7/NSP3-2A-S1f plasmids (Fig. 1). The total size of genome segments of the recombinant viruses, rSA11/3fS1, rSA11/S1f and rSA11/3fS1-His, is 20.7-20.8 Kbp, which is 2.1-2.2 Kbp (or ∼11%) greater than rSA11/wt, pointing to the ability of the rotavirus genome and virion particle to accommodate large amounts of foreign sequence.

**Figure 2.**
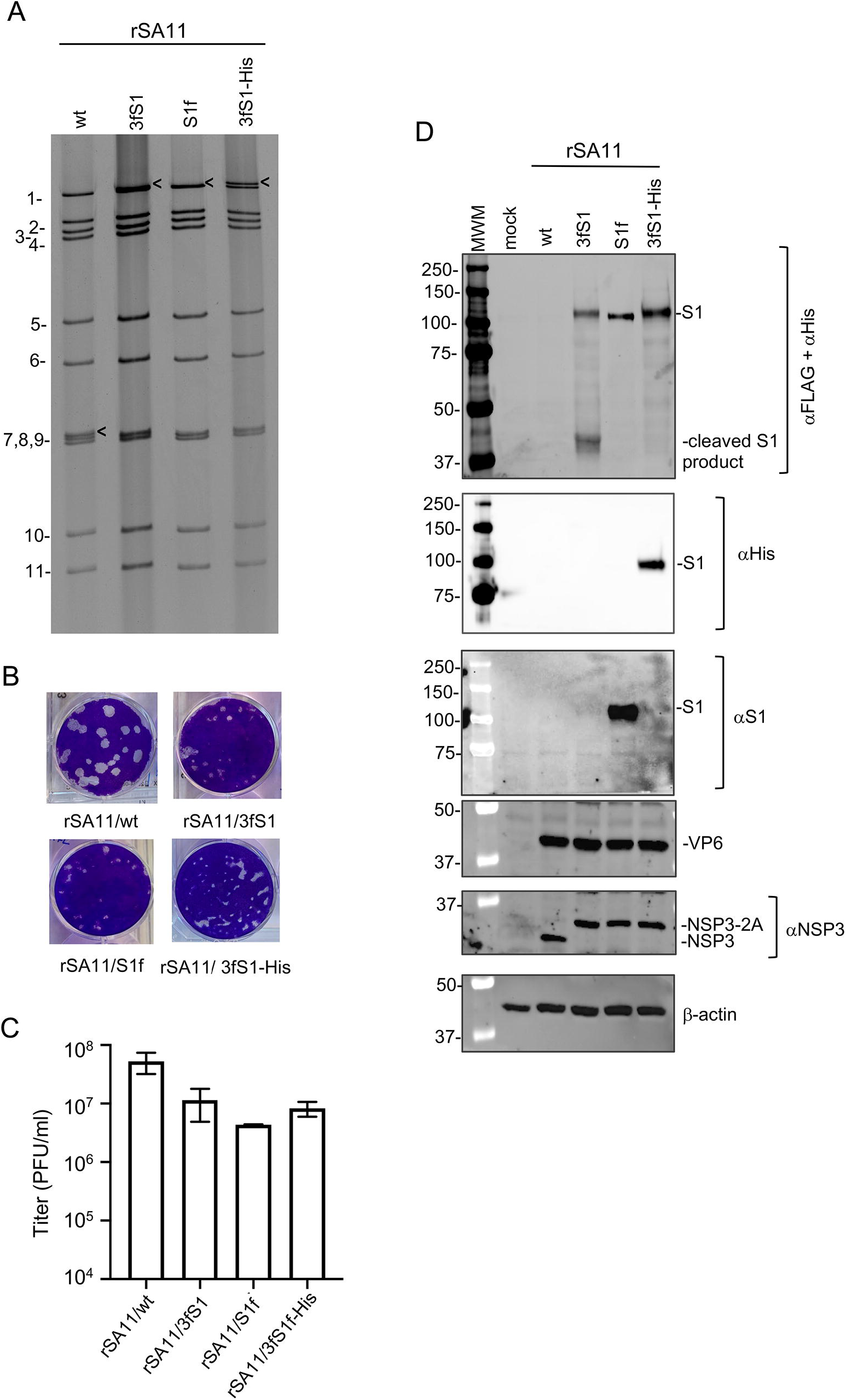
Properties of rSA11 viruses expressing SARS CoV-2 S1 proteins. **(A)** Viral dsRNA was recovered from MA104 cells infected with the indicated rSA11 isolates, resolved by gel electrophoresis, and detected by ethidium-bromide staining. RNA segments of rSA11/wt are labeled 1 to 11. Locations of segment 7 dsRNAs are identified with black arrows. **(B)** Plaque assays were performed using MA104 cells and detected by crystal-violet staining. **(C)** Titers reached by rSA11 isolates upon complete CPE in MA104 cells were determined by plaque assay. **(D)** Lysates were prepared from cells infected with the indicated rSA11 viruses at 9 h p.i and examined by immunoblot assay using FLAG/His antibodies to detect S proteins. The same blot was stripped and re-probed with antibodies specific for the SARS CoV-2 S1 (αS1) protein, rotavirus NSP3 and VP6, and cellular β-actin.

Analysis of the plaque phenotypes of rSA11/3fS1, rSA11/S1f and rSA11/3fS1-His showed that these viruses generated plaques on MA104 cell monolayers that were smaller than those produced by rSA11/wt (Fig. 2B). Similarly, recombinant viruses encoding the S1 protein grew to titers that were one-to two-logs lower than rSA11/wt (Fig. 2C). These findings are consistent with earlier reports which showed that the introduction of foreign sequences into segment 7 yields generates recombinant viruses with plaque phenotypes that are smaller in diameter than wild type virus and that grow to maximum titers that are less than wildtype virus (40, 42, 43). The poorer growth characteristics of recombinant viruses with larger segment 7 RNAs may stem from alterations in the quantity of NSP3+ produced in infected cells or in the translational efficiency of the NSP3+ RNA.

### rSA11 viruses expressing the S1 protein of SARS-CoV-2

To evaluate the nature of the S1 products expressed by recombinant viruses, lysates were prepared at 9 h p.i. from MA104 cells infected with rSA11/wt, rSA11/3fS1, rSA11/S1f and rSA11/3fS1-His. S1 products in the lysates were detected by immunoblot assay using anti-FLAG, anti-6xHis, and anti-S1 antibodies; rotavirus NSP3 and VP6 were detected using protein specific antisera. Analysis with anti-FLAG and anti-His antibodies (Fig. 2D) indicated that a ∼120 kD protein was expressed by rSA11/3fS1, rSA11/S1f and rSA11/3fS1-His but not by rSA11/wt. The 120-kD size corresponds to the glycosylated form of the S1 protein, suggesting that the S1 signal peptide in the recombinant viruses is functional. (The size of nonglycosylated S1 is 80kD). Production of the 120-kD protein by the modified segment 7 RNAs indicates that their 2A elements were functional enabling the expression of S1 as a separate protein product. Indeed, recombinant viruses with the modified segment 7 RNAs also produced an NSP3 product that was slightly larger in size than wild type NSP3 (Fig. 2D). The larger NSP3 results from the presence of 2A residues left behind at the C-terminus of NSP3 following 2A activity, suggesting that the 2A element was active in infected cells producing S1. Notably, immunoblot analysis indicated lysates from rSA11/NSP3-2A-3fS1-infected cells contained not only a ∼120-kD S1 product but also a small product of ∼40kD (Fig. 2D). The origin of the ∼40 kD product is unknown but may result from an aberrant cleavage of the 120-kD S1 protein. Interestingly, the ∼40kD product was not detected in lysates from cells infected with viruses expressing S1 with either a C-terminal FLAG or 6xHis tag, suggesting that modification of the C-terminus may affect the protein’s folding or modification such that the S1 protein becomes less prone to cleavage. Nonetheless, the instability of the S1 product may explain the poor detection of S1 expression reported earlier for the rSA11/3fS1 virus (43). Detection of the 120-kDa S1 proteins expressed by rSA11/S1f and rSA11/3fS1-His with antibodies recognizing tags present at the C-termini of the S1 proteins confirmed that the proteins represented full length products.

To further analyze the S1 products made by the rSA11/3fS1, rSA11/S1f and rSA11/3fS1-His viruses, we obtained a commercial rabbit polyclonal antibody (ABclonal A20136) generated against an S1 fusion protein (amino acids 11-682) (Fig. 2D). Unexpectedly, immunoblot analysis showed that the S1 antibody only recognized the 120-kDa product expressed by rSA11/S1f and not the 120-kDa products expressed by rSA11/3fS1 and rSA11/3fS1-His. The 120-kDa product of rSA11/S1f initiates with the S1 signal peptide, while the 120 kDa products of rSA11/3fS1 and rSA11/3fS1-His initiate with a 3xFLAG peptide, which is followed by the S1 signal peptide. It may be that the S1 antibody recognizes an epitope that is altered by the presence of an N-terminal tag and, as a result, the antibody cannot recognize the 120-kDa products of rSA11/3fS1 and rSA11/3fS1-His. Perhaps, for example, the N-terminal FLAG-tags of rSA11/3fS1 and rSA11/3fS1-His may affect glycosylation or folding of their S1 products in a manner that prevents recognition by the S1 antibody.

### Effect of endoglycosidase H (Endo H) treatment on expressed S1 protein

To further explore the possibility that the S1 products expressed by rSA11/3fS1, rSA11/S1f and rSA11/3fS1-His were glycosylated, we examined the effect that Endo H had on the electrophoretic migration patterns of the S1 products. To do this, we infected MA104 cells with the rSA11/3fS1, rSA11/S1f and rSA11/3fS1-His viruses, prepared lysates from the infected cells at 9 h p.i., and then treated portions of the lysates with Endo H using a Promega Endoglycosidase kit. The treated portions and untreated controls were resolved by gel electrophoresis, and immunoblot assay was used to determine the positions of the S1 products. Assays performed using the anti-FLAG and anti-6xHis antibodies showed that Endo H treatment significantly reduced the migration pattern of the S1 products of all three viruses from ∼120 kDa to 80-90 kDa (Fig. 3). These results indicate that the S1 products are heavily glycosylated and contain Endo H-sensitive asparagine-linked oligosaccharide chains. Immunoblot assays performed with the anti-S1 antibody likewise showed that Endo H caused a reduction in the size of the S1f product of the rSA11/S1f virus from ∼120 kDa to 80-90 kDa. Notably, the S1 antibody (ABclonal A20136) did not detect the Endo H-treated products of the rSA11/3fS1 and rSA11/3fS1-His viruses described above (Fig. 3). Endo H treatment truncates asparagine-linked oligosaccharide chains on proteins, leaving behind N-acetylglucosamine attached to asparagine. It may be possible that the presence of the residual N-acetylglucosamine residues interfere with the ability of the S1 antibody to recognize epitopes located in the S1 products of the rSA11/3fS1 and rSA11/3fS1-His viruses.

**Figure 3.**
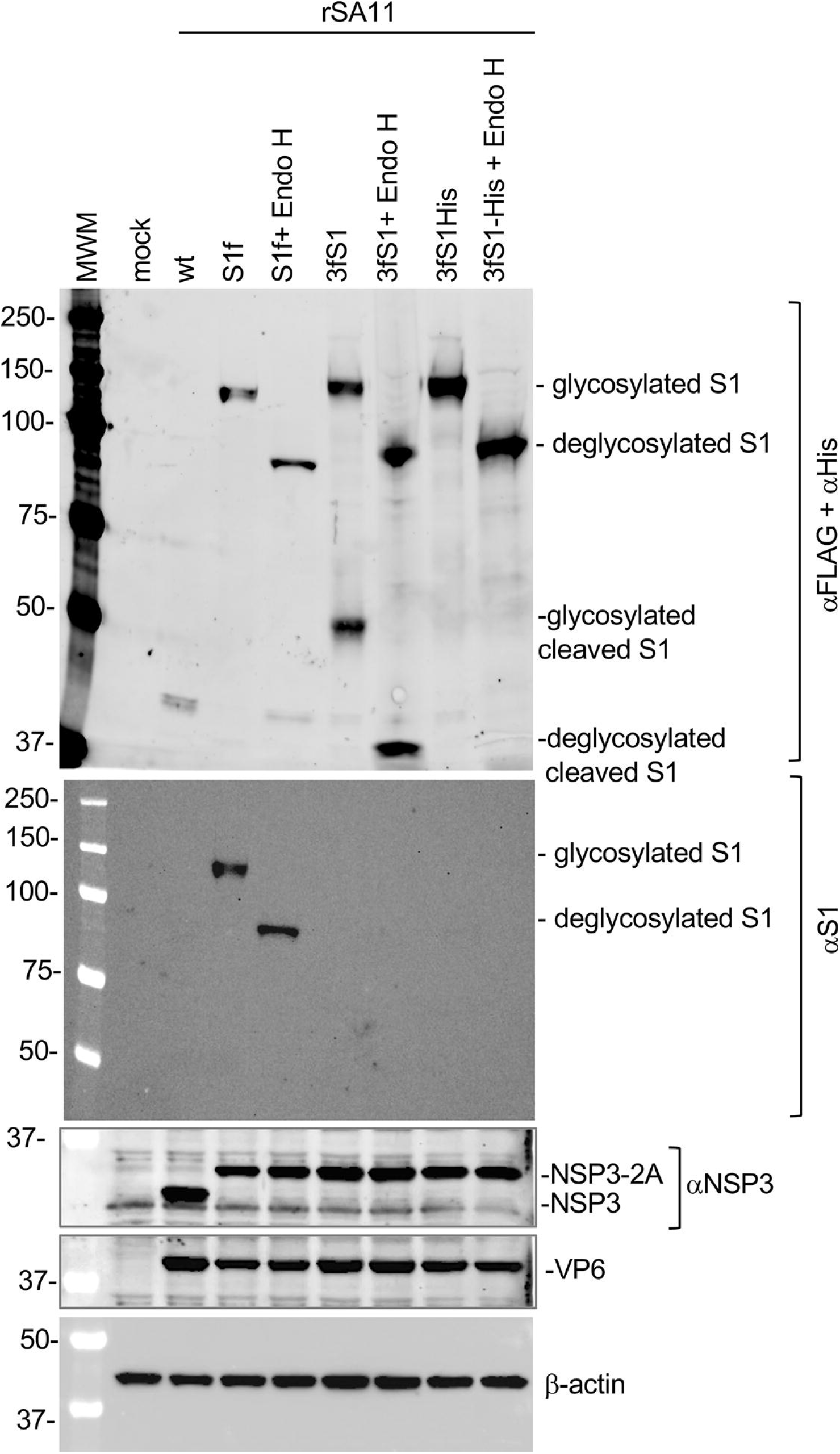
Endo H treatment of SARS CoV-2 S1 proteins expressed by rSA11 viruses. Whole cell lysates prepared at 9 h p.i. from MA104 cells infected with the indicated rSA11 viruses were mock-treated or treated with Endo H. The samples were examined by immunoblot assay using FLAG/His antibody to detect S1 proteins. The same blot was stripped and re-probed for (αS1) antibody specific for SARS CoV-2 S1. Immunoblots were also probed with antibodies specific for rotavirus NSP3 and VP6, and for β-actin.

### Binding of expressed S1 protein to the ACE2 receptor

Attachment of SARS-CoV-2 to target cells is mediated by the interaction of the RBD of the S protein with the human ACE2 (hACE2) transmembrane protein. To determine whether the S1 proteins expressed by rSA11 viruses were able to bind hACE2, we infected MA104 cells with rSA11/3fS1, rSA11/S1f and rSA11/3fS1-His. At 9 h p.i., clarified lysates prepared from the cells were incubated with soluble hACE2-Fc, a recombinant protein that contains the ACE2 extracellular domain fused to the Fc region of the IgG1 antibody. After incubation, protein A spin columns were used to recover the hACE2-Fc protein from the samples. After washing the spin columns, bound proteins were eluted and examined by immunoblot assay. Analysis of the eluted proteins by immunoblot analysis with anti-FLAG and anti-6xHis antibodies indicated that the S1 proteins expressed by rSA11/3fS1, rSA11/S1f and rSA11/3fS1-His viruses all bound hACE2-Fc (Fig. 4). This result suggests that the RBD domains of the S1 proteins generated by the rSA11 viruses are functional. Similar immunoblot analysis performed with anti-S1 antibody (A20136) also indicated that the S1 product of rSA11/S1f virus had affinity for hACE2-Fc and, thus, contained a functional RBD (Fig. 4). Because the S1 antibody does not recognize the S1 products of the rSA11/NSP3-2A-3fS1 and rSA11/NSP3-2A-3fS1-His viruses, it was not possible to use the S1 antibody in immunoblot assays to assess interaction of their S1 products with hACE2-Fc.

**Figure 4.**
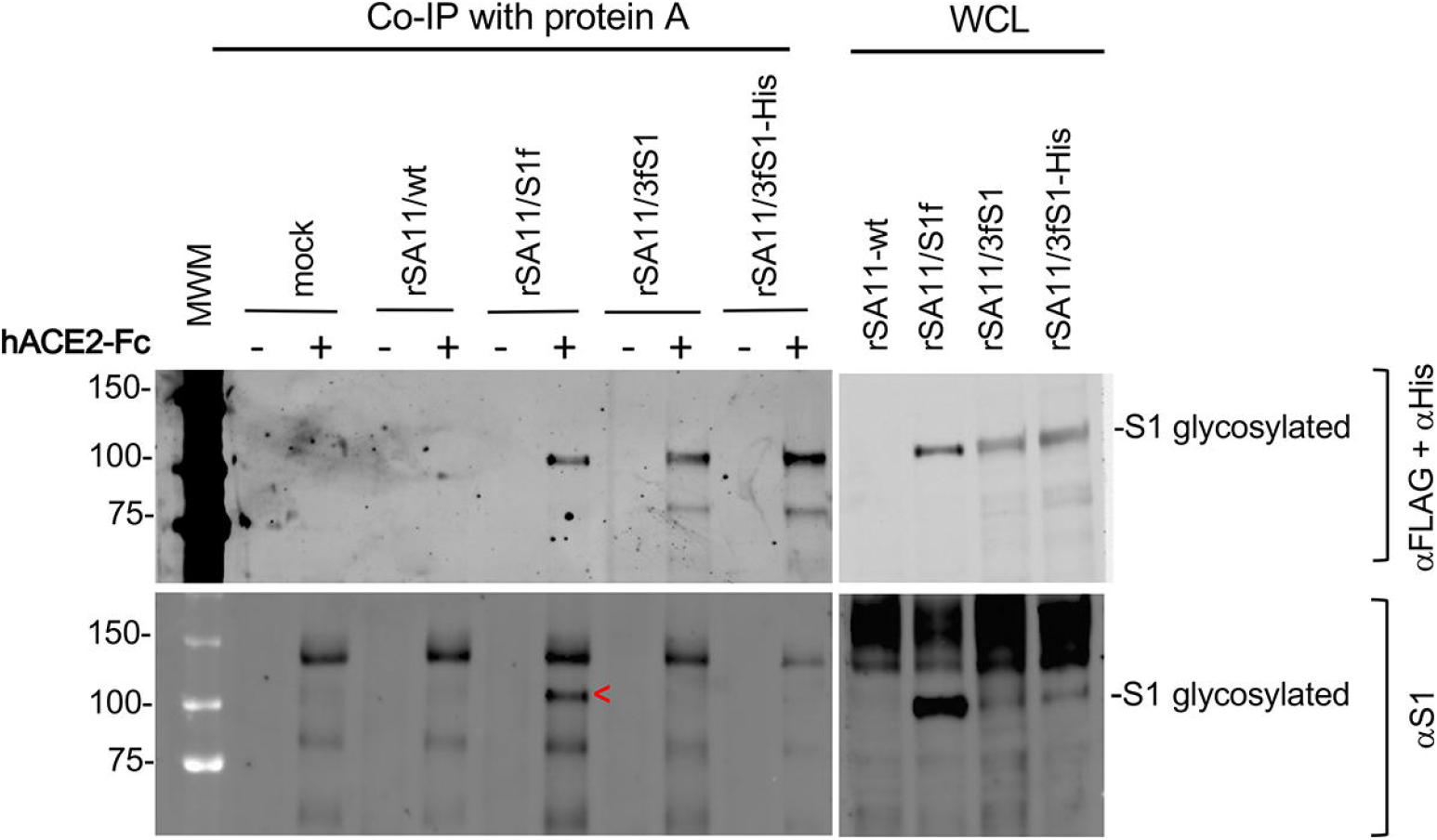
Binding of the SARS-CoV-2 S1 product to the human ACE2 receptor. Lysates prepared from MA104 cells infected with the indicated rSA11 viruses incubated with hACE2-Fc, a recombinant protein consisting of the ACE2 extracellular domain fused to Fc region of the IgG1 antibody Fc domain. Protein-A spin columns were used to recover hACE2-Fc, and associated proteins, from the lysates. The recovered samples were examined by immunoblot assay using antibodies specific for S1 products (FLAG/His antibody, SARS CoV-2 S1 antibody). MWM, molecular weight markers.

### Intracellular localization of expressed S1 protein

The results above indicated that only the S1f product of the rSA11/S1f virus contained a functional epitope for the anti-S1 antibody (A20136). To gain insight into the distribution of the S1f product in infected cells, MA104 cells were infected with rSA11/wt and rSA11/S1f viruses and then, at 9 h p.i., examined by IFA using the anti-S1 antibody, an antibody recognizing a major component of rotavirus replication factories (viroplasms, anti-NSP2), and an antibody recognizing a rotavirus ER/Golgi resident transmembrane protein (anti-NSP4). As expected, immunofluorescence signal was not detected in mock infected cells analyzed with the anti-S1, -NSP2, or -NSP4 antibodies (Fig. 5). Likewise, signal was not detected with the anti-S1 antibody in rSA11/wt-infected cells. Analysis of rSA11/S1f infected cells with the anti-S1 antibody indicated that a large portion of the S1f product accumulated in large cytoplasmic punctate structures (Fig. 5). The S1f fluorescence did not colocalize with viroplasms detected by the anti-NSP2 antibody, indicating the S1f product does not accumulate in rotavirus replication factories. In contrast, the S1f fluorescence co-colocalized extensively with the viral NSP4 ER/Golgi transmembrane protein (Fig. 5). This finding is consistent with results provided above that the S1f product is a N-mannose linked glycoprotein. Although the S1f product localizes to the same site of the infected cell where the outer capsid of the rotavirus particle is assembled (ER/Golgi) (2, 8), this does prevent the formation of infectious rotavirions (Fig. 2D). However, we cannot rule out the possibility the co-accumulation of S1f in the ER/Golgi may be partially responsible for the reduction of lower virus titers formed in rSA11/S1f-infected cells as compared to rSA11/wt-infected cells.

**Figure 5.**
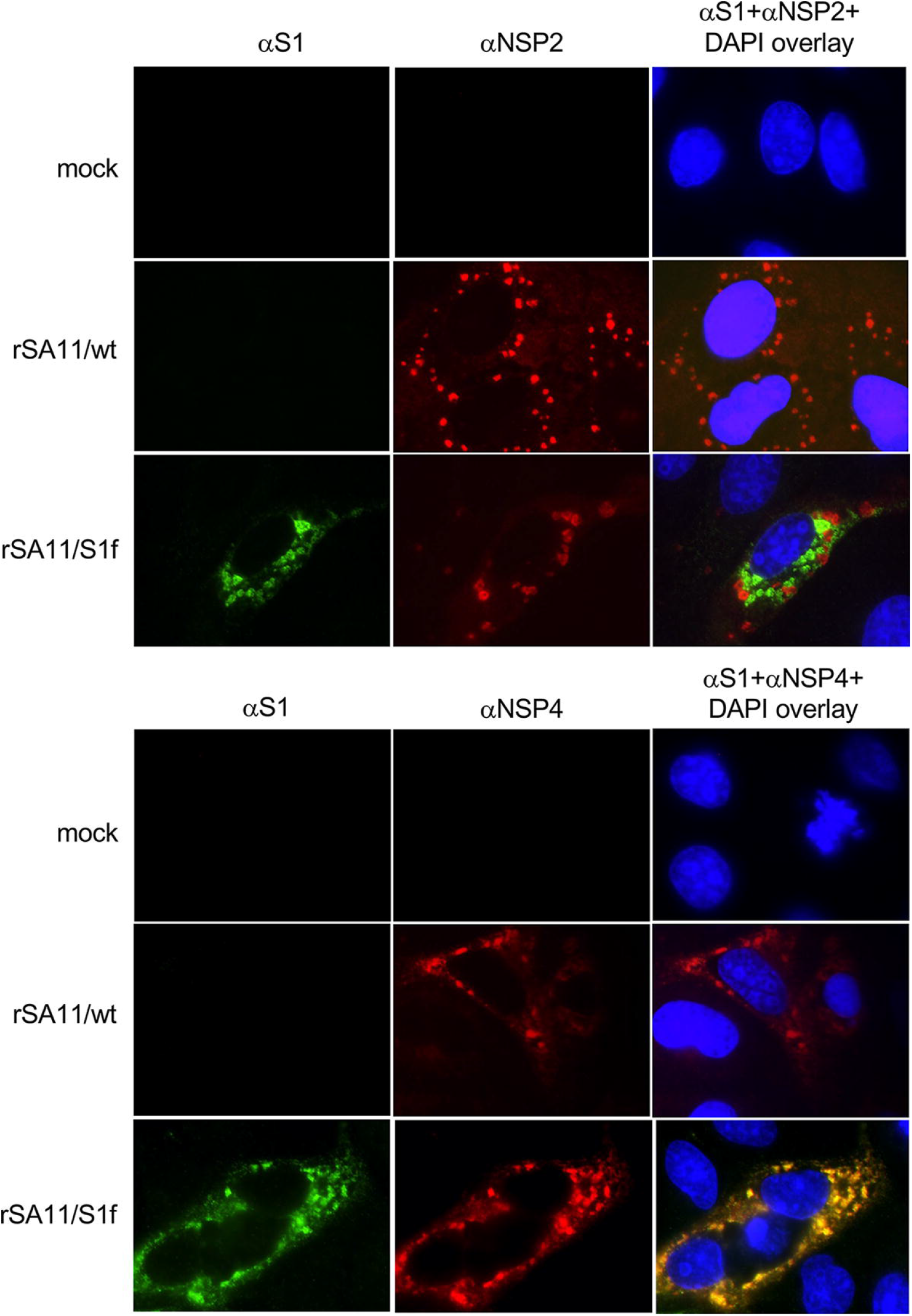
Localization of SARS CoV-2 S1 protein in rSA11/S1f-infected cells. MA104 cells were mock infected or infected with rSA11/S1f and, at 9 h p.i., fixed with ice cold methanol. Afterwards, the cells were incubated with rabbit S1 antibody, mouse NSP2 or NSP4 antibody, followed by Alexa 488 anti-rabbit IgG (green) and Alexa 594 anti-mouse IgG (red) to determine the locations of SARS-CoV-2 S1 and rotavirus NSP2 and NSP4 proteins. Nuclei were detected by staining with DAPI. Cells were analyzed with a Nikon Eclipse NiE microscope (100X oil immersion objective) and images were captured with a Hamamatsu Orca-Flash 2.8 sCMOS high resolution camera.

### Genetic stability of rSA11 viruses expressing S1f

Successful development of rotavirus as vaccine vector system requires the recombinant virus to be sufficiently genetically stable to allow scale up and production of large amounts of vaccine virus expressing a foreign sequence. To evaluate the genetic stability of rSA11/S1f, the virus was subjected to 5 rounds of serial passage at low multiplicity of infection in MA104 cells. In this procedure, infected cell lysates were diluted 1:10 to generate the inoculum used subsequent rounds of infection. Gel electrophoresis showed no difference in the viral dsRNA recovered from passage 1-to 5-infected cell lysates (Fig. 6), suggesting that the rSA11/S1f virus was genetically stable, at least though 5 rounds of serial passage. Immunoblot analysis showed that rSA11/S1f virus serially passaged up to 5 times in MA104 cells continued to direct the expression of the S1f product, consistent with the idea that this virus strain is genetically stable despite containing 2.1 Kbp of extra sequence. Interestingly, similar recombinant viruses (rSA11/ fVP1 and rSA11/VP1-h) carrying 1.7 Kbp of norovirus capsid sequence was genetically unstable, with its modified segment 7 fully lost from the virus pool upon 5 rounds of serial passage at 1:10 dilution. This finding suggests that it is not just the length of foreign sequence inserted into the rotavirus genome that influences genetic stability but also the nature of the foreign sequence (e.g., nucleotide composition, secondary structure) or the encoded protein.

**Figure. 6.**
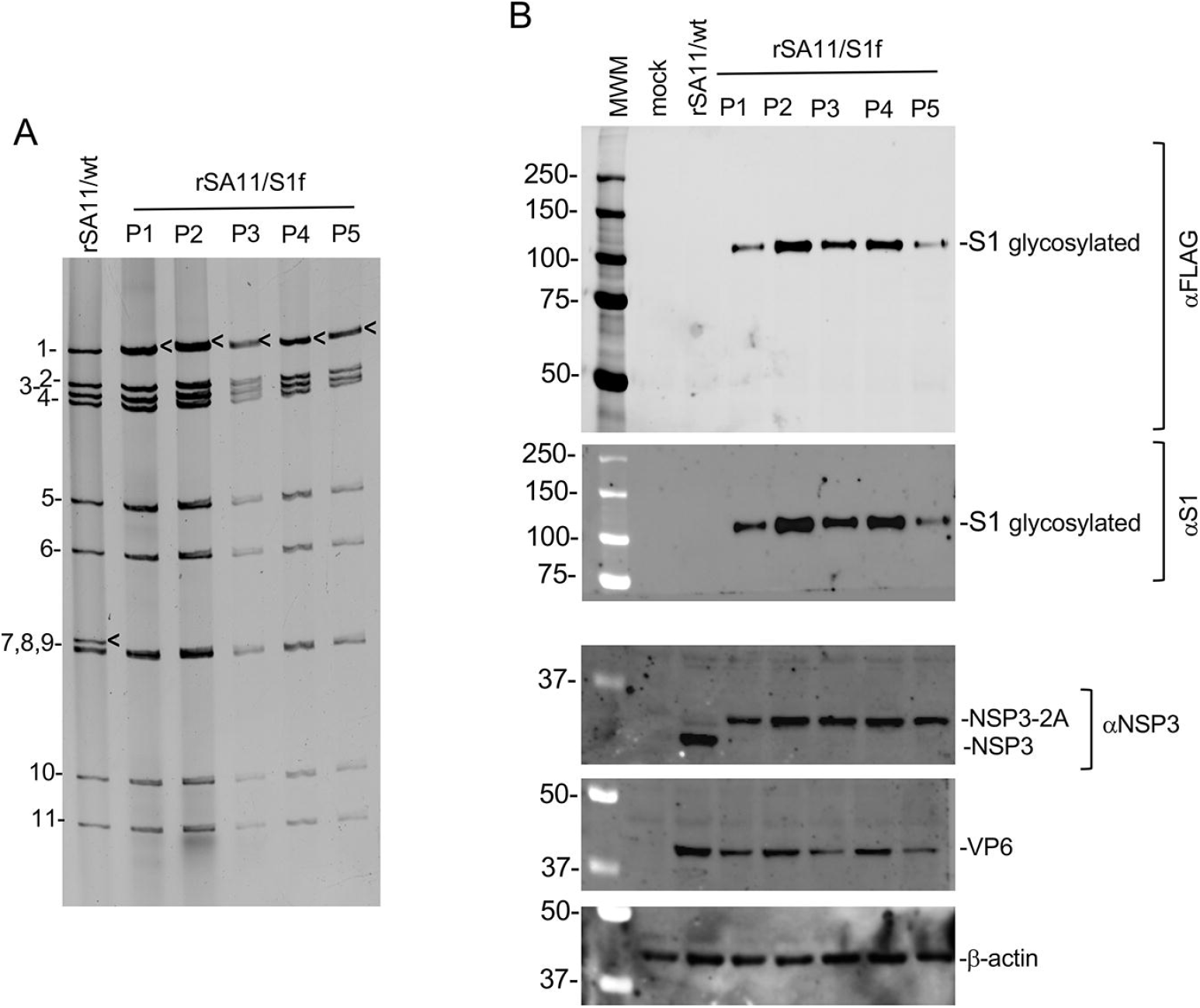
Genetic stability of rSA11 strains expressing SARS CoV-2 S1 f protein. The rSA11/S1f strain expressing S1 protein was serially passaged 5 times (P1 to P5) at 1:10 dilution in MA104 cells. **(A)** Genomic RNAs were recovered from infected cell lysates and analyzed by gel electrophoresis. Positions of viral genome segments are labeled. Position of modified segment 7 (NSP3) dsRNAs introduced into the rSA11 strain is denoted with black arrows. **(B)**. Lysates prepared from MA104 cells infected with rSA11/wt and serially passed SA11/NSP3-2A-S1f viruses (P1 to P5) were examined by immunoblot assay and probed with antibodies specific for FLAG, SARS-CoV-2 S1, rotavirus NSP3 and VP6, and β-actin. The top two panels were generated from the same blot. The αS1 blot (second panel) was produced after stripping the αFLAG blot (first panel) with WesternSure ECL stripping buffer (LI-COR).

## Discussion

In this study, we examined three rSA11 strains engineered to express the SARS-CoV-2 S1 through modification of the segment 7 (NSP3) dsRNA. Although the segment 7 RNAs of all three strains encoded S1 proteins with N-terminal signal peptides, two included sequence information that introduced a 3xFLAG tag immediately upstream of the signal peptide. The presence of the 3xFLAG was correlated with the inability of the S1 products of the rSA11/3fS1 and rSA11/3fS1-His viruses to be recognized by a widely used anti-S1 antibody (ABclonal A20136) (Fig. 2D). On the other hand, the S1f product of the rSA11/S1f virus, which lacks a peptide tag at is N-terminus, was readily detected by the anti-S1 antibody. The apparent size of the S1 product expressed by all three virus strains (rSA11/S1f, rSA11/3fS1, and rSA11/3fS1-His) was ∼120 kDa, which is consistent with glycosylated S1 generated during SARS-CoV-2 infection. Endo H treatment of the 120-kDa S1 product of the rSA11/S1f, rSA11/3fS1, and rSA11/3fS1-His viruses resulted in a size reduction to 80-90 kDa, providing additional evidence that the S1 products had undergone N-linked glycosylation (Fig. 3). Although all three S1 products were glycosylated, there were differences between the products that extended beyond the fact that only the S1f form could be recognized by the anti-S1 (A20136) antibody. For example, a significant portion of the 3fS1 product of the rSA11/3fS1 virus was cleaved in infected cells, yielding a major 40 kDa glycosylated S1 fragment (Fig. 2D). This cleavage was not seen for the S1f and 3fS1-His products of the rSA11/S1f and rSA11/3fS1-His viruses, suggesting that the 3fS1 has an aberrant protein-fold old that makes it prone to cleavage by an unidentified host protease. Collectively, our data indicates that the presence of a 3xFLAG tag upstream of the S1 signal peptide may impact the S1 product at multiple levels, including its glycosylation, fold structure and/or display of antibody epitopes. Our data also suggest that the S1f product of the rSA11/S1f virus has associated properties expected for authentic S1, suggesting that it may be possible to use rotaviruses as vaccine vectors, in developing a combined rotavirus-SARS-CoV-2 vaccine for immunization of infants and young children.

To our knowledge, this is the first study demonstrating that rotaviruses can be used as vector systems for the expression of glycosylated proteins. In previous work, we showed that by modifying the segment 7 dsRNA, it may be possible to develop rotaviruses as vaccine vectors for the expression of nonglycosylated capsid proteins of other enteric and mucosal viruses, such as norovirus (42), hepatitis A, C, and E viruses, polioviruses, adenoviruses. and enteroviruses. The results here show that it may be possible to extend the use of rotaviruses to include their development as vaccine vectors for the expression of the glycosylated capsid proteins of other enteric and mucosal viruses (HIV, coronavirus, Zika virus, and Ebola virus). The widespread use of rotavirus vaccines throughout the world makes rotavirus vectors prime candidates for the development of combination vaccines in infants and young children.

Our findings confirm previous observations that the rotavirus genome can accommodate a remarkable amount of foreign sequence (40, 42, 43). Based on work with the rSA11/S1 virus strains (43, and herein), the rotavirus genome can accommodate at least 2.1 Kbp of additional RNA sequence. A surprising finding from this study was that the genetic stability of rSA11/S1 viruses was remarkably greater than what has been observed for other rSA11 strains, in which smaller amounts of foreign sequence had been introduced into the rotavirus genome (42). Discovering what factors influence genetic stability will be important for fully developing rotaviruses as vaccine vectors systems.

## Acknowledgement

We appreciate the support and suggestions of the members of the IU family of virologists on this project. We are also indebted to Dr. Ulla Buchholz (Laboratory of Infectious Diseases, NIID, NIH) for the gift of BHK-T7 cells.

## Contributions

Conceptualization, JTP; Methodology, JTP, SH and AAP; Investigation, SH and AAP; Formal analysis, JTP and AAP; Writing—original draft preparation, JTP; Writing—review and editing, JTP and AAP; Supervision, JTP; Funding acquisition, JTP. All authors have read and agreed to the published version of the manuscript

## References

1. Matthijnssens J, Attoui H, Bányai K, Brussaard CPD, Danthi P, Del Vas M, Dermody TS, Duncan R, Fāng !方勤 Q, Johne R, Mertens PPC, Mohd Jaafar F, Patton JT, Sasaya 笹谷孝英 T, Suzuki 鈴木信弘 N, Wei 魏太云 T. ICTV Virus Taxonomy Profile: *Sedoreoviridae* 2022. J Gen Virol. 2022 Oct;103(10).

2. Crawford SE, Ramani S, Tate JE, Parashar UD, Svensson L, Hagbom M, Franco MA, Greenberg HB, O’Ryan M, Kang G, Desselberger U, Estes MK. 2017. Rotavirus infection. Nat Rev Dis Primers 3: 17083.

3. Clark A, Black R, Tate J, Roose A, Kotloff K, Lam D, Blackwelder W, Parashar U, Lanata C, Kang G, Troeger C, Platts-Mills J, Mokdad A; Global Rotavirus Surveillance Network, Sanderson C, Lamberti L, Levine M, Santosham M, Steele D. 2017. Estimating global, regional and national rotavirus deaths in children aged <5 years: Current approaches, new analyses and proposed improvements. PLoS One 12:e0183392.

4. Leshem E, Tate JE, Steiner CA, Curns AT, Lopman BA, Parashar UD. 2015. Acute gastroenteritis hospitalizations among US children following implementation of the rotavirus vaccine. JAMA 313:2282–2284; Erratum in 2015, 314:188.

5. Burke RM, Tate JE, Kirkwood CD, Steele AD, Parashar UD. 2019. Current and new rotavirus vaccines. Curr Opin Infect Dis 32: 435–444.

6. Folorunso OS, Sebolai OM. 2020. Overview of the development, impacts, and challenges of live-attenuated oral rotavirus vaccines. Vaccines (Basel) 8(3): 341.

7. Skansberg A, Sauer M, Tan M, Santosham M, Jennings MC. 2021. Product review of the rotavirus vaccines ROTASIIL, ROTAVAC, and Rotavin-M1. Hum Vaccin Immunother 17:1223-1234.

8. Trask SD, McDonald SM, Patton JT. 2012. Structural insights into the coupling of virion assembly and rotavirus replication. Nat Rev Microbiol 10:165–177

9. Desselberger U. 2020. What are the limits of the packaging capacity for genomic RNA in the cores of rotaviruses and of other members of the Reoviridae? Virus Res 276:197822.

10. Kanai Y, Komoto S, Kawagishi T, Nouda R, Nagasawa N, Onishi M, Matsuura Y, Taniguchi K, Kobayashi T. 2017. Entirely plasmid-based reverse genetics system for rotaviruses. Proc Natl Acad Sci USA 114: 2349–2354.

11. Komoto S, Fukuda S, Kugita M, Hatazawa R, Koyama C, Katayama K, Murata T, Taniguchi K. 2019. Generation of infectious recombinant human rotaviruses from just 11 cloned cDNAs encoding the rotavirus genome. J Virol 93(8): e02207–18.

12. Kanda M, Fukuda S, Hamada N, Nishiyama S, Masatani T, Fujii Y, Izumi F, Okajima M, Taniguchi K, Sugiyama M, Komoto S, Ito N. 2022. Establishment of a reverse genetics system for avian rotavirus A strain PO-13. J Gen Virol 103(6), 10.1099.

13. Sánchez-Tacuba L, Feng N, Meade NJ, Mellits KH, Jaïs PH, Yasukawa LL, Resch TK, Jiang B, López S, Ding S, Greenberg HB. 2020. An optimized reverse genetics system suitable for efficient recovery of simian, human, and murine-like rotaviruses. J Virol 94:e01294–20.

14. Komoto, S., Kanai, Y., Fukuda, S., Kugita, M., Kawagishi, T., Ito, N., Sugiyama, M., Matsuura, Y., Kobayashi, T., Taniguchi, K. 2017. Reverse genetics system demonstrates that rotavirus nonstructural protein NSP6 is not essential for viral replication in cell culture. J Virol 91 pii:e00695-17.

15. Chang-Graham AL, Perry JL, Strtak AC, Ramachandran NK, Criglar JM, Philip AA, Patton JT, Estes MK, Hyser JM. 2019. Rotavirus calcium dysregulation manifests as dynamic calcium signaling in the cytoplasm and endoplasmic reticulum. Sci Rep 9:10822.

16. Papa G, Venditti L, Arnoldi F, Schraner EM, Potgieter C, Borodavka A, Eichwald C, Burrone OR. 2019. Recombinant rotaviruses rescued by reverse genetics reveal the role of NSP5 hyperphosphorylation in the assembly of viral factories. J Virol 94: e01110–e01119.

17. Falkenhagen A, Patzina-Mehling C, Gadicherla AK, Strydom A, O’Neill HG, Johne R. 2020. Generation of simian rotavirus reassortants with VP4-and VP7-encoding genome segments from human strains circulating in Africa using reverse genetics. Viruses 12:201.

18. Falkenhagen A, Patzina-Mehling C, Rückner A, Vahlenkamp TW, Johne R. 2019. Generation of simian rotavirus reassortants with diverse VP4 genes using reverse genetics. J Gen Virol 100:1595–1604.

19. Criglar JM, Crawford SE, Zhao B, Smith HG, Stossi F, Estes MK. 2020. A genetically engineered rotavirus NSP2 phosphorylation mutant impaired in viroplasm formation and replication shows an early interaction between vNSP2 and cellular lipid droplets. J Virol 94, e00972–20.

20. Song Y, Feng N, Sanchez-Tacuba L, Yasukawa LL, Ren L, Silverman RH, Ding S, Greenberg HB. 2020. Reverse genetics reveals a role of rotavirus VP3 phosphodiesterase activity in inhibiting RNase L signaling and contributing to intestinal viral replication *in vivo*. J Virol 94:e01952-19.

21. Nilsson EM, Sullivan OM, Anderson ML, Argobright HM, Shue TM, Fedowitz FR, LaConte LEW, Esstman SM. 2021. Reverse genetic engineering of simian rotaviruses with temperature-sensitive lesions in VP1, VP2, and VP6. Virus Res 302:198488.

22. Hundley F, Biryahwaho B, Gow M, Desselberger U. 1985. Genome rearrangements of bovine rotavirus after serial passage at high multiplicity of infection. Virology 143:88–103.

23. Hundley F, McIntyre M, Clark B, Beards G, Wood D, Chrystie I, Desselberger U. 1987. Heterogeneity of genome rearrangements in rotaviruses isolated from a chronically infected immunodeficient child. J Virol 61:3365–3372.

24. Shen S, Burke B, Desselberger U. 1994. Rearrangement of the VP6 gene of a group A rotavirus in combination with a point mutation affecting trimer stability. J Virol 68:1682–1688.

25. Ballard A, McCrae MA, Desselberger U 1992. Nucleotide sequences of normal and rearranged RNA segments 10 of human rotaviruses. J Gen Virol 73:633–638.

26. Gault E, Schnepf N, Poncet D, Servant A, Teran S, Garbarg-Chenon A. 2001. A human rotavirus with rearranged genes 7 and 11 encodes a modified NSP3 protein and suggests an additional mechanism for gene rearrangement. J Virol 75:7305–7314.

27. Patton JT, Taraporewala Z, Chen D, Chizhikov V, Jones M, Elhelu A, Collins M, Kearney K, Wagner M, Hoshino Y, Gouvea V. 2001. Effect of intragenic rearrangement and changes in the 3’ consensus sequence on NSP1 expression and rotavirus replication. J Virol 75:2076–2086.

28. Barro M, Patton JT. 2005. Rotavirus nonstructural protein 1 subverts innate immune response by inducing degradation of IFN regulatory factor 3. Proc Natl Acad Sci USA 102:4114–4119.

29. Morelli M, Dennis AF, Patton JT. 2015. Putative E3 ubiquitin ligase of human rotavirus inhibits NF-κB activation by using molecular mimicry to target β-TrCP. mBio 6:e02490–14.

30. Arnold MM, Brownback CS, Taraporewala ZF, Patton JT. 2012. Rotavirus variant replicates efficiently although encoding an aberrant NSP3 that fails to induce nuclear localization of poly(A)-binding protein. J Gen Virol 93:1483–1494.

31. Gratia M, Sarot E, Vende P, Charpilienne A, Baron CH, Duarte M, Pyronnet S, Poncet D. 2015. Rotavirus NSP3 is a translational surrogate of the poly(A)-binding protein-poly(A) complex. J Virol 89: 8773–8782.

32. Piron M, Delaunay T, Grosclaude J, Poncet D. 1999. Identification of the RNA-binding, dimerization, and eIF4GI-binding domains of rotavirus nonstructural protein NSP3. J Virol 73:5411–5421.

33. Hatazawa R, Fukuda S, Kumamoto K, Matsushita F, Nagao S, Murata T, Taniguchi K, Matsui T, Komoto S. 2021. Strategy for generation of replication-competent recombinant rotaviruses expressing multiple foreign genes. J Gen Virol. 102(4).

34. Komoto S, Fukuda S, Ide T, Ito N, Sugiyama M, Yoshikawa T, Murata T, Taniguchi K. 2018. Generation of recombinant rotaviruses expressing fluorescent proteins by using an optimized reverse genetics system. J Virol 92: e00588–18.

35. Kanai Y, Kawagishi T, Nouda R, Onishi M, Pannacha P, Nurdin JA, Nomura K, Matsuura, Y, Kobayashi T. 2018. Development of stable rotavirus reporter expression systems. J Virol 93:e01774–18.

36. Pannacha P, Kanai Y, Kawagishi T, Nouda R, Nurdin JA, Yamasaki M, Nomura K, Lusiany T, Kobayashi T. 2021. Generation of recombinant rotaviruses encoding a split NanoLuc peptide tag. Biochem Biophys Res Commun 534:740–746.

37. Guanghui Y, Dai J, Patton JT. 2021. Contributions of RING finger domain and C-terminus of rotavirus NSP1 to efficient viral replication. (in review).

38. Philip AA, Herrin BE, Garcia ML, Abad AT, Katen SP, Patton JT. 2019. Collection of recombinant rotaviruses expressing fluorescent reporter proteins. Microbio Resour Announc 8(27): e00523–19.

39. Philip AA, Perry JL, Eaton HE, Shmulevitz M, Hyser JM, Patton JT. 2019. Generation of recombinant rotavirus expressing NSP3-UnaG fusion protein by a simplified reverse genetics system. J Virol 93: e01616–19.

40. Philip AA, Patton JT. 2020b. Expression of separate heterologous proteins from the rotavirus NSP3 genome segment using a translational 2A stop-restart element. J Virol 94: e00959–20.

41. Luke G, Escuin H, De Felipe P, Ryan M. 2020. 2A to the fore - research, technology and applications. Biotechnol Genet Eng Rev 26:223–260.

42. Philip AA, Patton JT. 2022. Generation of Recombinant Rotaviruses Expressing Human Norovirus Capsid Proteins. J Virol 96:e0126222.

43. Philip AA, Patton JT. 2021. Rotavirus as an expression platform of domains of the SARS-CoV-2 spike protein. Vaccines (Basel) 9:449.

44. Duan L, Zheng Q, Zhang H, Niu Y, Lou Y, Wang H. 2020. The SARS-CoV-2 spike glycoprotein biosynthesis, structure, function, and antigenicity: implications for the design of spike-based vaccine immunogens. Front Immunol 11:576622.

45. Huang Y, Yang C, Xu XF, Xu W, Liu SW. 2020. Structural and functional properties of SARS-CoV-2 spike protein: potential antivirus drug development for COVID-19. Acta Pharmacol Sin 41:1141–1149.

46. Brouwer PJM, Caniels TG, van der Straten K, Snitselaar JL, Aldon Y, Bangaru S, Torres JL, Okba NMA, Claireaux M, Kerster G, et al. 2020. Potent neutralizing antibodies from COVID-19 patients define multiple targets of vulnerability. Science 369:643–650.

47. Liu L, Wang P, Nair MS, Yu J, Rapp M, Wang Q, Luo Y, Chan JF, Sahi V, Figueroa A, et al. 2020. Potent neutralizing antibodies against multiple epitopes on SARS-CoV-2 spike. Nature 584:450–456.

48. Rogers TF, Zhao F, Huang D, Beutler N, Burns A, He WT, Limbo O, Smith C, Song G, Woehl J, Yang L, Abbott RK, Callaghan S, Garcia E, Hurtado J, Parren M, Peng L, Ramirez S, Ricketts J, Ricciardi MJ, Rawlings SA, Wu NC, Yuan M, Smith DM, Nemazee D, Teijaro JR, Voss JE, Wilson IA, Andrabi R, Briney B, Landais E, Sok D, Jardine JG, Burton DR. 2020. Isolation of potent SARS-CoV-2 neutralizing antibodies and protection from disease in a small animal model. Science 369:956–963.

49. Zost SJ, Gilchuk P, Chen RE, Case JB, Reidy JX, Trivette A, Nargi RS, Sutton RE, Suryadevara N, Chen EC, et al. 2020. Rapid isolation and profiling of a diverse panel of human monoclonal antibodies targeting the SARS-CoV-2 spike protein. Nat Med 26:1422– 1427.

50. Xiaojie S, Yu L, Lei Y, Guang Y, Min Q. 2021. Neutralizing antibodies targeting SARS-CoV-2 spike protein. Stem cell research 50:102125.

51. Casalino L, Gaieb Z, Goldsmith JA, Hjorth CK, Dommer AC, Harbison AM, Fogarty CA, Barros EP, Taylor BC, McLellan JS, Fadda E, Amaro RE. 2020. Beyond shielding: the roles of glycans in the SARS-CoV-2 spike protein. ACS Cent Sci 6:1722–1734.

52. Medina-Enríquez MM, Lopez-León S, Carlos-Escalante JA, Aponte-Torres Z, Cuapio A, Wegman-Ostrosky T. 2020. ACE2: the molecular doorway to SARS-CoV-2. Cell Biosci 10:148.

53. Philip AA, Dai J, Katen SP, Patton JT. 2020. Simplified reverse genetics method to recover recombinant rotaviruses expressing reporter proteins. J Vis Exp 158: e61039.

54. Arnold M, Patton JT, McDonald SM. 2009. Culturing, storage, and quantification of rotaviruses. Curr Protoc Microbiol Chapter 15:Unit 15C.3.

